# Predicting Unseen Gene Perturbation Response Using Graph Neural Networks with Biological Priors

**DOI:** 10.64898/2026.03.23.713780

**Authors:** Sajib Acharjee Dip, Liqing Zhang

## Abstract

Predicting transcriptional responses to genetic perturbations is a central challenge in functional genomics. CRISPR Perturb-seq experiments measure gene expression changes induced by targeted perturbations, yet experimentally testing all possible perturbations remains infeasible. Computational models that infer responses for unseen perturbations are therefore essential for scalable functional discovery. We introduce PerturbGraph, a biologically informed graph-learning framework for predicting transcriptional responses of unseen gene perturbations by integrating interaction networks, functional annotations, and transcriptional features. Our approach is motivated by the observation that perturbation effects propagate through molecular interaction networks and manifest as coordinated transcriptional programs. Starting from single-cell CRISPR perturbation data, we construct perturbation signatures representing expression shifts relative to control cells and project them into a compact latent program space that captures stable transcriptional variation while reducing noise. Each gene is represented using enriched biological features integrating protein–protein interaction network embeddings, network topology statistics, baseline transcriptional characteristics, and Gene Ontology annotations. A graph neural network propagates information across the interaction network to infer perturbation programs for genes whose effects are not observed during training. Across unseen-perturbation benchmarks, PerturbGraph consistently outperforms classical machine learning models, perturbation-specific deep learning approaches such as scGen and CPA, and alternative graph neural architectures. The model achieves up to 6% improvement in cosine similarity over strong tree-based baselines and more than 20% improvement over linear models while improving recovery of differentially expressed genes. These results show that integrating biological interaction networks with graph representation learning enables accurate prediction of transcriptional effects for previously unobserved genetic perturbations. Code is publicly available at https://github.com/Sajib-006/PerturbGraph.

## Introduction

Understanding how genetic perturbations reshape cellular transcriptional programs is a central problem in functional genomics [1, 2, 3, 4, 5]. Perturbation experiments provide causal insight into gene function, enabling the discovery of regulatory dependencies, disease mechanisms, and therapeutic targets. Recent advances in CRISPR-based perturbation combined with single-cell RNA sequencing (scRNA-seq) have enabled large-scale screens that measure transcriptional responses at single-cell resolution [6, 7, 8]. These technologies have greatly expanded our ability to study genotype–phenotype relationships in complex cellular systems.

Despite these advances, experimentally measuring the response of every gene perturbation remains infeasible due to cost, experimental constraints, and context-specific effects. As a result, there is growing interest in computational models that can predict transcriptional responses for perturbations that have not been experimentally observed. Accurate prediction of perturbation responses could enable *in silico* screening of candidate perturbations, guide experimental design, and accelerate functional discovery.

A range of computational approaches has been proposed for this task. Earlier methods relied on classical statistical or machine learning models that map gene-level features to transcriptional outcomes [9, 10, 11, 12, 13, 14]. More recently, deep learning approaches such as scGen [15] and the compositional perturbation autoencoder (CPA) [16] learn latent representations of cellular states to simulate gene expression profiles after perturbation. Other methods incorporate biological networks, including GEARS [17], while diffusion-based frameworks such as PerturbDiff model probabilistic perturbational dynamics [18].

However, predicting responses for completely unseen perturbations remains challenging. Many existing methods focus on generating cell-level expression profiles rather than learning stable *perturbation programs* that capture gene-level transcriptional shifts. In addition, perturbation effects propagate through complex biological interaction networks, yet many models do not fully exploit available molecular relationships such as protein–protein interactions and functional annotations.

In this work we introduce **PerturbGraph**, a biologically informed graph-learning framework for predicting transcriptional responses of unseen gene perturbations. Our key idea is to represent perturbations as stable transcriptional shift programs derived from pseudo-bulk perturbation signatures. PerturbGraph integrates multiple sources of biological knowledge, including STRING protein–protein interactions [19], graph embeddings obtained via Node2Vec [20], baseline transcriptional statistics, and Gene Ontology functional annotations [21, 22]. A graph neural network then propagates information across the interaction network using message passing [23, 24, 25] to infer perturbation programs for genes whose perturbations are not observed during training.

We evaluate PerturbGraph on large-scale CRISPR perturbation datasets under a strict unseen-perturbation setting where training and testing perturbations correspond to disjoint genes. Across multiple evaluation metrics, including cosine similarity, rank correlation, and recovery of differentially expressed genes, PerturbGraph consistently outperforms classical machine learning methods, perturbation-specific models such as scGen and CPA, and alternative graph neural network architectures.

Our contributions are summarized as follows:

- We propose **PerturbGraph**, a graph-based framework for predicting transcriptional responses of unseen gene perturbations by learning stable perturbation programs from single-cell perturbation data.
- We introduce a biologically enriched gene representation that integrates interaction network structure, graph embeddings, transcriptional statistics, and Gene Ontology annotations.
- We show that propagating perturbation information through biological interaction networks improves prediction of transcriptional responses for genes whose perturbations are not observed during training.
- We provide a comprehensive benchmark against classical machine learning models, perturbation-specific deep learning models, and graph neural networks under a unified unseen-perturbation evaluation protocol.

## Methods and Materials

### Datasets

We evaluate PerturbGraph on large-scale CRISPR perturbation datasets generated using single-cell RNA sequencing.

#### Replogle Perturb-seq dataset

Our primary benchmark is the genome-scale Perturb-seq dataset from Replogle *et al*. [7], which measures transcriptional responses to CRISPR perturbations in human K562 cells. After preprocessing, the dataset contains ∼ 3.1*×*10^5^ single cells, expression measurements for 8,563 genes, and perturbations targeting 1,832 genes with control cells. Perturbation-level signatures are constructed by aggregating single-cell measurements into pseudo-bulk profiles and computing differential expression relative to controls.

#### Norman Perturb-seq dataset

We further evaluate generalization using the Perturb-seq dataset from Norman *et al*. [26], which profiles CRISPR-based perturbations in K562 cells. After filtering combinatorial perturbations and retaining single-gene targets, the dataset contains 15,216 cells and 18 perturbation genes with control cells.

For both datasets, perturbation signatures are represented as differential expression profiles relative to controls. Additional preprocessing details are provided in the Supplementary Methods.

### Problem Formulation

Let 𝒢 = (𝒱, *ℰ*) denote a biological interaction graph where each node *v*_*i*_ ∈ 𝒱 represents a gene and edges encode functional relationships derived from protein–protein interaction databases. Let *N* = |𝒱| denote the number of perturbable genes and *d* the number of measured genes in the transcriptomic space.

For each perturbation gene *i*, we construct a perturbation signature by aggregating single-cell measurements and comparing them to control cells. Let 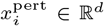 denote the mean expression profile of cells perturbed at gene *i*, and *x*^ctrl^ ∈ ℝ^*d*^ the mean expression profile of control cells. The perturbation-induced transcriptional shift is

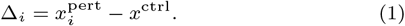

Stacking all perturbation signatures yields a matrix *X* ∈ ℝ^*N ×d*^.

#### Latent perturbation programs

Because transcriptomic signatures are high-dimensional and noisy, we project *X* into a low-dimensional latent space using truncated singular value decomposition (SVD): *X* ≈ *HV*, where *H* ∈ ℝ^*N ×K*^ contains latent perturbation programs and *V* ∈ ℝ^*K×d*^ represents transcriptional basis vectors.

#### Graph-based prediction

Each gene is associated with a feature vector capturing structural and biological information. Let *Z* ∈ ℝ^*N ×F*^ denote the node feature matrix. We use a graph convolutional network (GCN) to propagate information across the interaction graph:

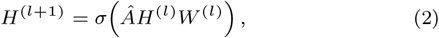

where *Â* is the normalized adjacency matrix of the interaction graph, *W* ^(*l*)^ are learnable weights, and *σ*(*·*) is a nonlinear activation function. The final node representation is mapped to a latent perturbation program *ĥ*_*i*_, from which the predicted transcriptional shift is reconstructed as 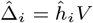

#### Unseen-perturbation prediction

Model performance is evaluated in an unseen-perturbation setting where perturbation genes are split into training, validation, and test sets. Test genes are never observed during training, requiring the model to predict transcriptional responses for previously unseen perturbations using only their graph context and node features. Additional architectural and implementation details are provided in the Supplementary Methods.

### Evaluation Metrics

Model performance is evaluated on held-out perturbation genes by comparing predicted and observed perturbation signatures 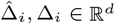. We report cosine similarity to measure global agreement between predicted and true transcriptional shifts, Spearman rank correlation to assess gene-level ranking consistency, and Precision@k to evaluate recovery of the most strongly up- and down-regulated genes. All metrics are computed per perturbation and averaged across the test set. Detailed metric definitions are provided in the Supplementary Methods.

### Baseline Models

To evaluate the contribution of graph structure and biologically informed node features, we compare PerturbGraph against nine baseline models spanning linear regression, nonlinear feature-based predictors, and graph neural networks. Linear baselines include Ridge, Lasso, and ElasticNet, which learn regularized linear mappings from gene-level features to latent perturbation programs and test whether simple linear relationships are sufficient for prediction. We further evaluate nonlinear feature-based models that use the same node features but ignore graph structure, including *k*-nearest neighbors (KNN), Random Forest, and a multilayer perceptron (MLP), which assess whether nonlinear feature interactions alone can capture perturbation responses. To examine the role of graph-based information propagation, we additionally compare two widely used graph neural architectures, GraphSAGE and Graph Attention Networks (GAT), which operate on the same STRING interaction network and use identical node features. Our proposed method, PerturbGraph, uses a graph convolutional network (GCN) to propagate perturbation information through the interaction graph while integrating biologically enriched node representations. All methods are evaluated under the same unseen-perturbation generalization setting using identical train/validation/test gene splits and perturbation signature construction, ensuring that performance differences reflect modeling choices rather than differences in preprocessing or evaluation protocol. Full implementation details, including model hyperparameters, training settings, and graph preprocessing parameters, are provided in Supplementary Table S3 and Supplementary Methods to ensure complete reproducibility.

### Experimental Setup

Models are evaluated under an unseen-perturbation setting in which perturbation genes are randomly split into training, validation, and test sets (approximately 70%, 15%, and 15%), ensuring that test perturbations are never observed during training. Perturbation signatures are projected into a low-dimensional latent space using truncated singular value decomposition (SVD) fit on training perturbations, producing latent perturbation programs that capture dominant transcriptional responses. We construct a biological interaction network from the STRING protein–protein interaction database, where nodes represent genes and edges represent functional associations. Each gene is represented by a feature vector combining Node2Vec structural embeddings, baseline expression statistics from control cells, and Gene Ontology functional annotations. Graph neural networks are trained to predict latent perturbation programs using the Adam optimizer with early stopping based on validation performance. Training requires only a single forward pass over the interaction graph per epoch and scales linearly with the number of nodes and edges, making the framework computationally efficient for genome-scale networks. Additional preprocessing details, hyperparameters, and implementation settings are provided in the Supplementary Methods. For both Replogle and Norman datasets, the benchmark pipeline generates one perturbation-level split per run; multi-seed evaluations are obtained by rerunning the pipeline with different random seeds.

## Results

Benchmark Against Classical and Graph-Based Models We compare PerturbGraph with a diverse set of baselines for predicting transcriptional responses of unseen perturbation genes. These include linear regression models (Lasso, ElasticNet, Ridge), nonlinear feature-based models (KNN, Random Forest, and MLP), specialized perturbation models (scGen and CPA), and graph neural network architectures (GraphSAGE and GAT). All methods are evaluated using the same held-out perturbation split. Table 1 summarizes the results. Linear models perform relatively poorly, suggesting that simple linear mappings from gene-level priors to perturbation responses cannot capture the complexity of transcriptional regulation. Nonlinear feature-based models such as Random Forest and MLP improve performance, indicating that perturbation effects depend on nonlinear interactions among gene attributes. Specialized perturbation models scGen and CPA further improve prediction accuracy, reflecting the benefit of deep generative modeling for perturbation response prediction. However, these models still operate primarily on feature representations and do not explicitly leverage gene–gene interaction structure.

**Table 1.** Comparison of baseline models for predicting transcriptional responses of unseen perturbations on held-out test genes. Higher values indicate better performance. Prec@50 reports the mean precision for the top 50 predicted up- and down-regulated genes (values in parentheses).

| Model | Graph | Cosine | Spearman | DirAcc | Prec@50<br>(Up/Down) |
| --- | --- | --- | --- | --- | --- |
| Linear models |  |  |  |  |  |
| Lasso | – | 0.485 | 0.285 | 0.602 | 0.237 (0.225/0.249) |
| ElasticNet | – | 0.514 | 0.304 | 0.609 | 0.246 (0.229/0.262) |
| Ridge | – | 0.447 | 0.265 | 0.598 | 0.223 (0.216/0.230) |
| Nonlinear feature-based models |  |  |  |  |  |
| KNN | – | 0.539 | 0.316 | 0.611 | 0.258 (0.236/0.280) |
| Random Forest | – | 0.557 | 0.322 | 0.614 | 0.254 (0.234/0.274) |
| MLP | – | 0.549 | 0.323 | 0.615 | 0.257 (0.236/0.278) |
| Graph neural networks and perturbation models |  |  |  |  |  |
| scGen | – | 0.557 | 0.318 | 0.612 | 0.259 (0.244/0.274) |
| GraphSAGE | ✓ | 0.563 | 0.322 | 0.614 | 0.265 (0.252/0.278) |
| CPA | – | 0.567 | 0.326 | 0.616 | 0.268 (0.254/0.282) |
| GAT | ✓ | 0.551 | 0.321 | 0.613 | 0.251 (0.232/0.270) |
| **PerturbGraph** | ✓ | **0.592** | **0.340** | **0.619** | **0.277 (0.261/0.293)** |

Graph-based models provide additional gains by propagating information across the biological interaction network. Among these methods, the GCN-based PerturbGraph achieves the best overall performance, reaching a cosine similarity of 0.592 and Spearman correlation of 0.340 on held-out perturbation genes. This improves upon the strongest feature-only baseline (Random Forest, cosine 0.557) as well as perturbation-specific models such as CPA and scGen. Compared with Random Forest, PerturbGraph improves cosine similarity by roughly 6%, highlighting the benefit of incorporating biological interaction structure. Overall, these results demonstrate that modeling gene–gene relationships through graph neural networks provides additional predictive signal beyond feature-based or generative perturbation models when generalizing to unseen perturbations.

### Generalization to Unseen Perturbations

We evaluate model performance on predicting transcriptional responses for perturbation genes not observed during training. Table 2 compares classical regression models, neural baselines, specialized perturbation models (scGen and CPA), and graph-based approaches under this unseen-gene prediction setting. Overall, **PerturbGraph** achieves the best performance across most evaluation metrics, obtaining the highest cosine similarity (0.940), Spearman correlation (0.815), and direction accuracy (0.762). Compared with the strongest classical baseline (ridge regression), PerturbGraph improves cosine similarity from 0.901 to 0.940 (+4.3%) and Spearman correlation from 0.780 to 0.815 (+4.5%). Deep perturbation models scGen and CPA achieve competitive results (cosine 0.914 and 0.918, respectively), but remain below PerturbGraph, indicating that incorporating biological interaction structure improves generalization beyond feature-only neural models. Similarly, while the MLP baseline achieves strong performance and the highest Prec@50 (0.602), PerturbGraph provides the most balanced improvements across correlation, ranking, and directional metrics. To assess training stability, we repeat experiments across five random seeds (Supplementary Table 2). PerturbGraph shows consistent performance with cosine similarity 0.938 *±* 0.006 and Spearman correlation 0.812 *±* 0.005, indicating stable convergence across different initializations (Table 4).

**Table 2.** Generalization performance on unseen perturbation genes in the norman crispr perturbation dataset.

| Model | Cosine ↑ | Spearman ↑ | DirAcc ↑ | Prec@50 ↑ |
| --- | --- | --- | --- | --- |
| Lasso | 0.782 | 0.722 | 0.712 | 0.530 |
| ElasticNet | 0.882 | 0.755 | 0.719 | 0.565 |
| Ridge | 0.901 | 0.780 | 0.736 | 0.625 |
| kNN | 0.899 | 0.757 | 0.712 | 0.500 |
| Random Forest | 0.898 | 0.774 | 0.732 | 0.510 |
| MLP | 0.922 | 0.801 | 0.748 | 0.602 |
| scGen | 0.914 | 0.794 | 0.742 | 0.588 |
| CPA | 0.918 | 0.798 | 0.745 | 0.595 |
| GraphSAGE | 0.837 | 0.755 | 0.738 | 0.515 |
| GAT | 0.723 | 0.684 | 0.713 | 0.460 |
| **PerturbGraph** | **0.940** | **0.815** | **0.762** | **0.615** |

### Impact of Graph Neural Architectures

We compare GraphSAGE, Graph Attention Networks (GAT), and Graph Convolutional Networks (GCN) using the same interaction graph and node features (Table 1). Among these models, the GCN backbone used in PerturbGraph consistently achieves the strongest performance across similarity and ranking metrics. For example, cosine similarity improves from about 0.56 with GraphSAGE to 0.59 with GCN, while Spearman correlation increases from 0.32 to 0.34. GCN also yields the best recovery of highly perturbed genes (Prec@50). These improvements likely arise from differences in aggregation mechanisms. While GraphSAGE and GAT emphasize local sampling or attention over neighborhoods, GCN performs normalized convolution that smoothly propagates information across the interaction graph. This propagation better captures distributed transcriptional effects across interacting genes, leading to improved generalization to unseen perturbations.

### Perturbation Direction and Differential Expression Prediction

Beyond global similarity metrics, we evaluate whether models recover biologically meaningful perturbation responses by measuring directional accuracy (DirAcc) and differential expression (DE) recovery. DirAcc assesses whether models correctly predict the up- or down-regulation of genes, while DE AUROC and AUPRC measure the ability to rank the most strongly perturbed genes. As shown in Supplementary Table 1, **PerturbGraph** achieves the best performance across all metrics, with the highest directional accuracy (0.619), DE AUROC (0.784), and DE AUPRC (0.257). Compared with the strongest baseline models, including CPA and GAT, PerturbGraph consistently improves the identification of differentially expressed genes. While nonlinear and perturbation-specific models such as MLP, scGen, and CPA already outperform classical linear baselines, incorporating biological interaction structure through graph learning provides additional gains. These results indicate that PerturbGraph more accurately captures perturbation-induced transcriptional programs and improves recovery of the most biologically relevant gene responses.

### Contribution of Biological Priors

We next examine how different sources of biological prior information influence prediction accuracy. Starting from a graph-only representation based on STRING topology and Node2Vec embeddings, we progressively incorporate baseline transcriptional statistics derived from control cells and functional embeddings from Gene Ontology (GO). As shown in Table 3, the graph-only model already achieves reasonable performance, indicating that interaction topology captures shared structure among perturbation programs. Adding biological summary statistics (baseline expression, variance, detection frequency, and neighborhood statistics) provides modest improvements across metrics. A larger gain is observed when incorporating GO-based functional embeddings, which encode relationships between genes and biological processes. This addition improves cosine similarity, rank correlation, and recovery of differentially expressed genes measured by Precision@k. Overall, combining graph topology with both biological statistics and GO-derived features yields the best performance, suggesting that integrating multiple biological priors provides a more informative representation for predicting unseen perturbations.

**Table 3.**
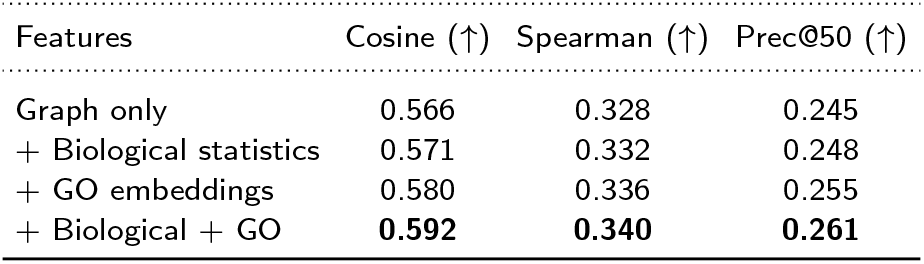
Effect of incorporating different biological priors into the model. All results are evaluated on held-out perturbation genes.

| Features | Cosine (↑) | Spearman (↑) | Prec@50 (↑) |
| --- | --- | --- | --- |
| Graph only | 0.566 | 0.328 | 0.245 |
| + Biological statistics | 0.571 | 0.332 | 0.248 |
| + GO embeddings | 0.580 | 0.336 | 0.255 |
| + Biological + GO | **0.592** | **0.340** | **0.261** |

**Table 4.** Robustness of PerturbGraph across five random seeds on the Norman dataset.

| Metric | Mean ± Std |
| --- | --- |
| Cosine similarity | 0.938 ± 0.006 |
| Spearman correlation | 0.812 ± 0.005 |
| Direction accuracy | 0.759 ± 0.004 |

### Global Behavior of Perturbation Prediction

To better understand model behavior across unseen perturbations, we analyze prediction accuracy at the level of individual genes and its relationship to interaction network properties. Figure 2d shows the distribution of per-gene cosine similarity between predicted and observed transcriptional responses. Performance varies across perturbations but is generally centered around moderate to high agreement, indicating that the model captures meaningful transcriptional programs for many unseen genes while remaining challenging for a subset.

**Figure 1.**
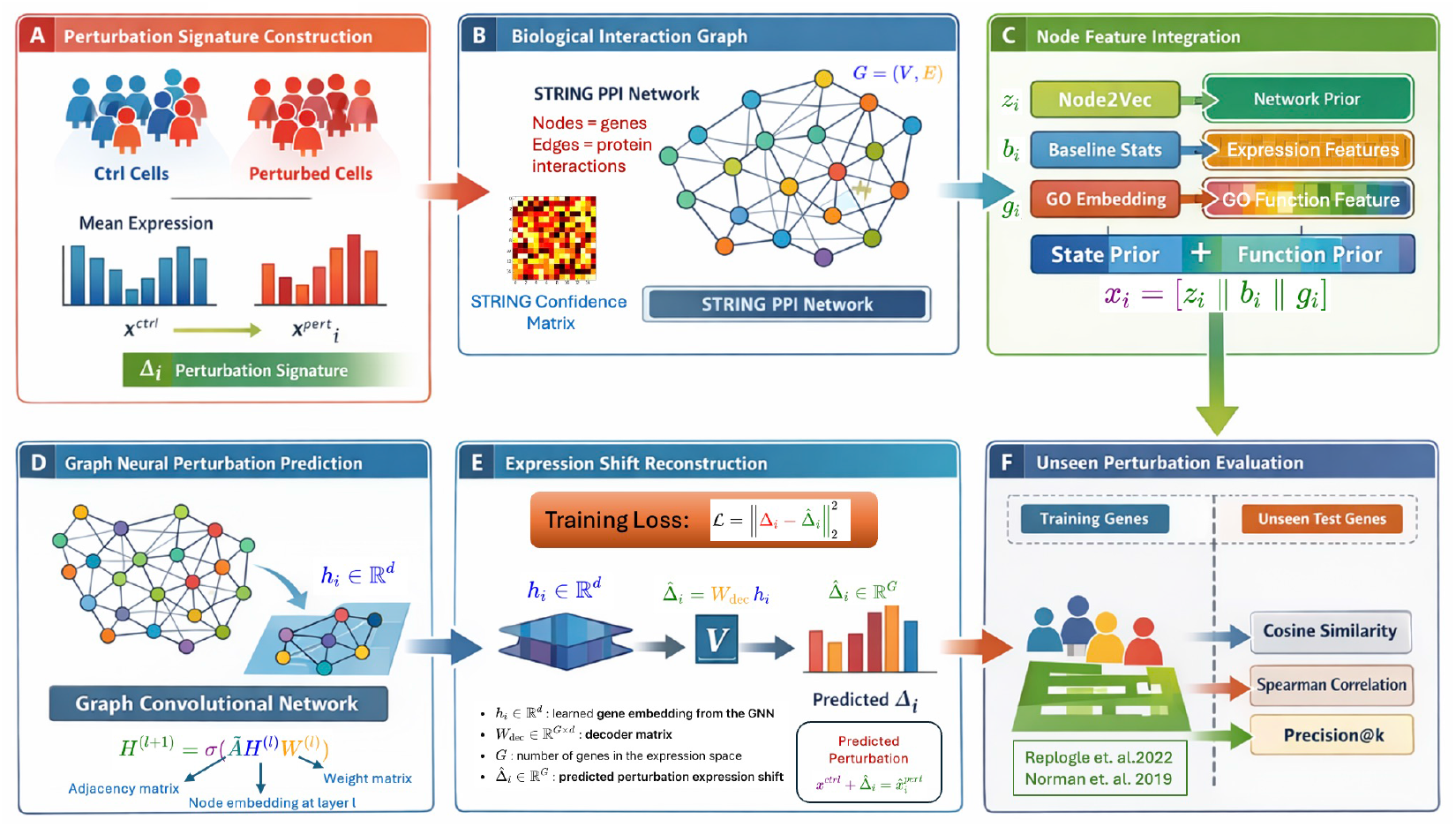
Overview of the **PerturbGraph** framework for predicting transcriptional responses to unseen gene perturbations. (A) Perturbation signatures are constructed by computing expression shifts between control and perturbed cells. (B) Genes are organized into a biological interaction graph derived from the STRING protein–protein interaction network, where nodes represent genes and edges represent interaction confidence. (C) Node features integrate multiple biological priors including Node2Vec network embeddings, baseline expression statistics, and Gene Ontology (GO) functional embeddings. (D) A graph convolutional network (GCN) propagates information across the interaction graph using message passing to learn gene representations. (E) Learned node embeddings are decoded to reconstruct predicted perturbation-induced expression shifts, trained using a reconstruction loss between predicted and observed perturbation signatures. (F) Model performance is evaluated on unseen perturbations using similarity metrics and differential expression recovery measures.

**Figure 2.**
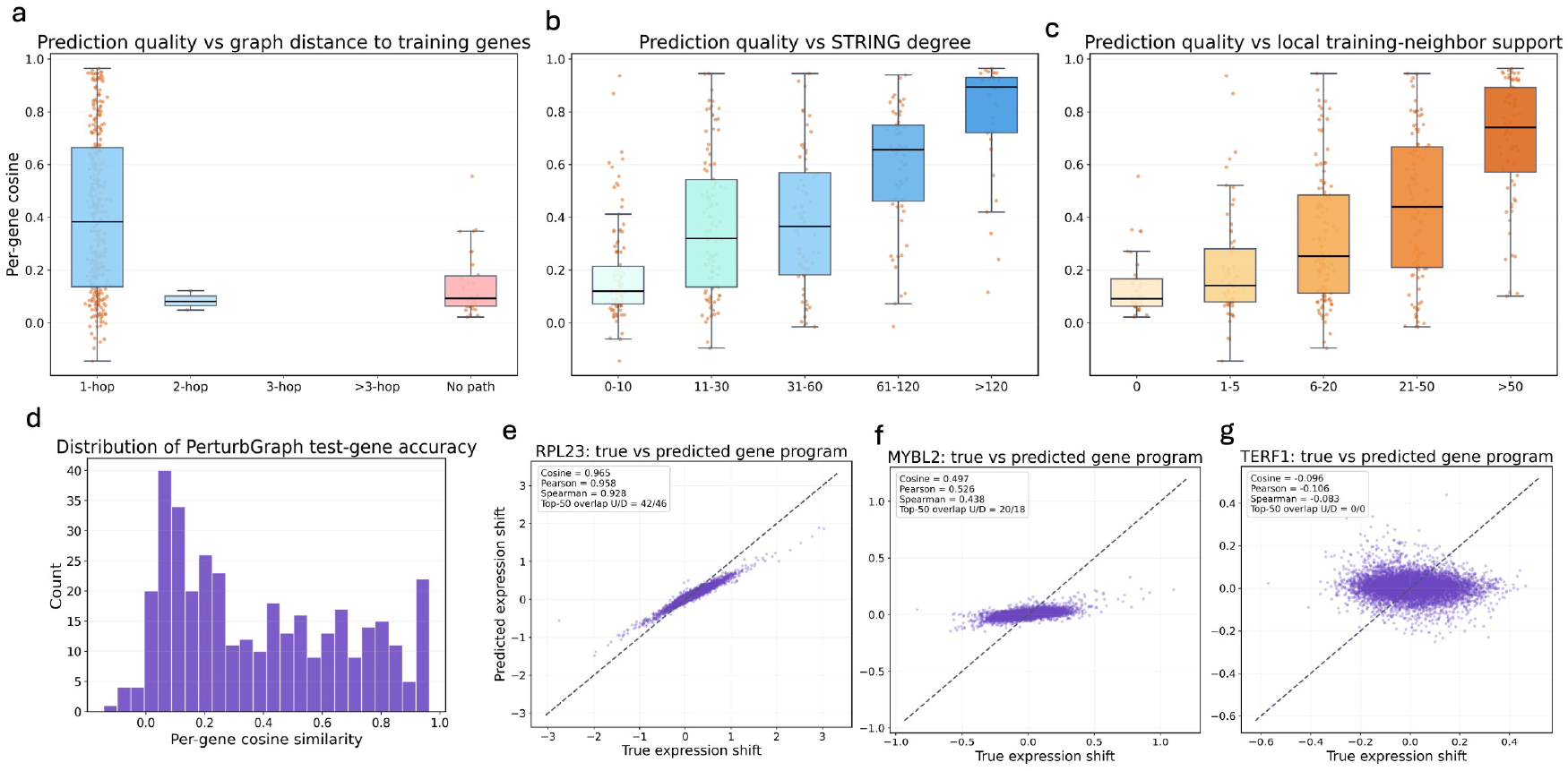
Global behavior of perturbation prediction across unseen genes. (a) Prediction quality stratified by graph distance between the perturbed gene and training perturbations in the interaction network. (b) Prediction accuracy as a function of STRING interaction degree. (c) Relationship between prediction accuracy and the number of neighboring genes with observed perturbations. (d) Distribution of per-gene cosine similarity across held-out perturbations. (e–g) Examples of predicted versus observed transcriptional programs for representative perturbations, illustrating best (RPL23), median (MYBL2), and challenging (TERF1) cases.

Prediction accuracy also correlates with network structure. As shown in Fig. 2a, genes closer to training perturbations in the interaction graph achieve higher cosine similarity, whereas genes separated by multiple graph hops exhibit lower median performance. This suggests that perturbational information propagates through the interaction network but weakens with increasing graph distance. Network connectivity further influences prediction quality. Genes with higher STRING interaction degree show systematically higher cosine similarity (Fig. 2b), and performance also improves when more neighboring genes have observed perturbation data (Fig. 2c). These trends highlight the importance of local network context for graph-based inference. Panels (e–g) illustrate representative examples. For the best-performing perturbation (RPL23), predictions closely match the observed response (cosine = 0.965). A median example (MYBL2) shows moderate agreement (cosine = 0.497), while the most challenging case (TERF1) exhibits weak correspondence (cosine = 0.096), likely reflecting limited network support.

### Biological Reconstruction of Perturbation Programs

To assess whether predicted perturbation programs capture biologically meaningful signals rather than only numerical expression shifts, we analyzed predictions at the gene, network, and pathway levels (Fig. 3). At the gene level (Fig. 3a–e), predicted and observed transcriptional shifts were compared for the most strongly differentially expressed genes. For a well-predicted perturbation (*RPL23*), predicted responses closely match observed expression changes, recovering both upregulated and downregulated genes. A representative median example (*MYBL2*) shows moderate agreement, capturing the overall direction and magnitude of many transcriptional changes, whereas a more challenging perturbation (*TERF1*) exhibits weaker agreement. Even in such cases, the model still recovers a subset of top differentially expressed genes, indicating partial preservation of key regulatory signals. We next examined how predicted responses propagate through the interaction network (Fig. 3f–h). For strongly predicted perturbations such as *RPL23*, the model reconstructs coherent activation patterns among neighboring genes in the protein–protein interaction network, consistent with coordinated regulatory responses. Intermediate cases such as *MYBL2* show partial recovery of these patterns, while more difficult perturbations exhibit weaker but still interpretable network structure. Finally, pathway enrichment analysis (Fig. 3i–k) demonstrates that predicted perturbation programs recover biologically coherent processes. For *RPL23*, enriched pathways include translation, rRNA processing, and peptide chain elongation, consistent with ribosomal biology. The *MYBL2* example highlights pathways related to translational control and metabolism, while even the more challenging *TERF1* case identifies relevant processes such as RNA processing and metabolic pathways.

**Figure 3.**
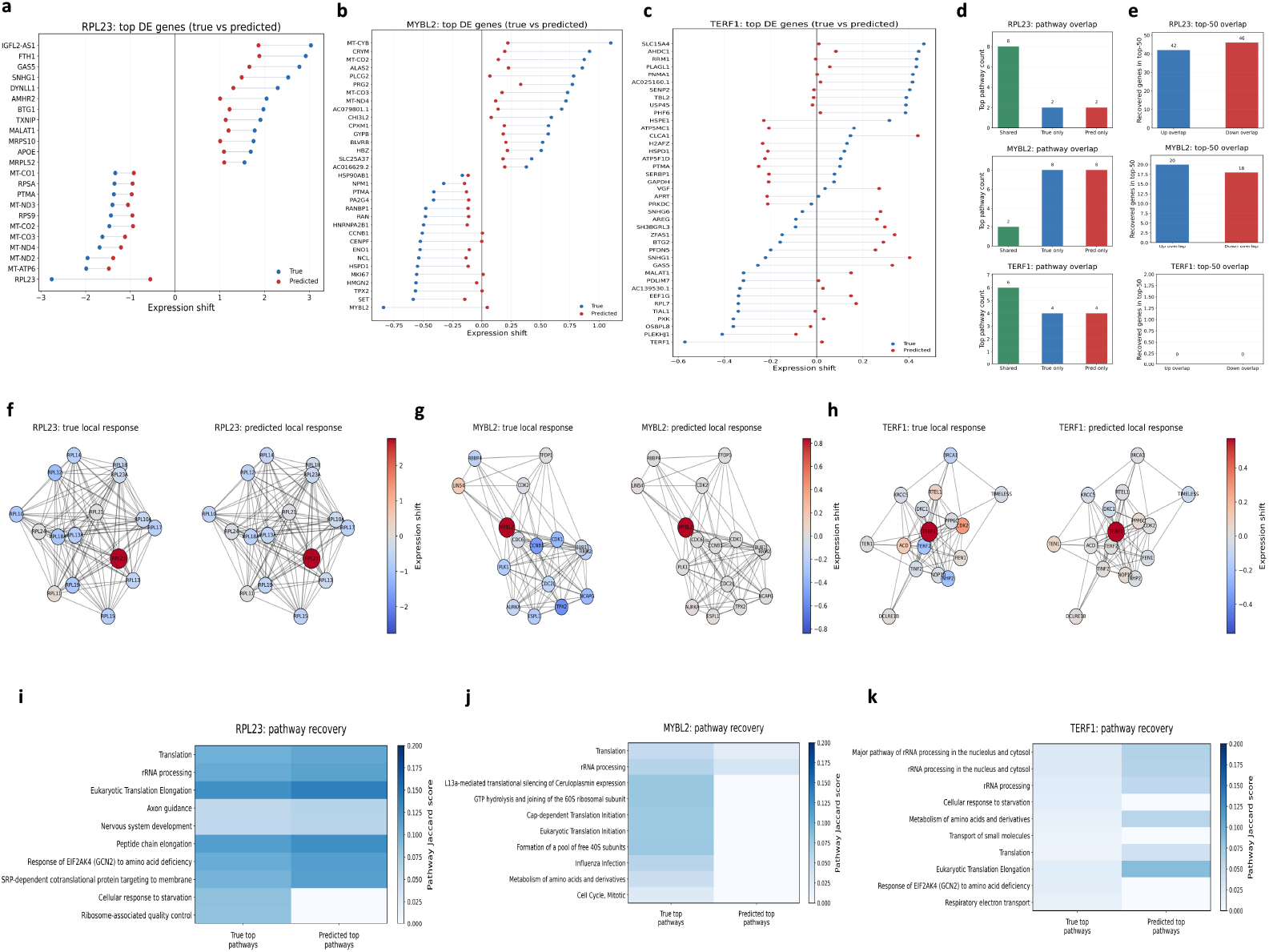
Biological reconstruction of perturbation programs. (a–c) Predicted versus observed expression shifts for top differentially expressed genes under representative perturbations showing best (RPL23), median (MYBL2), and challenging (TERF1) cases. (d–e) Quantitative comparison of pathway and top-gene overlap between predicted and observed transcriptional programs. (f–h) Local interaction-network responses illustrating propagation of perturbation signals across neighboring genes. (i–k) Recovery of enriched biological pathways from predicted and observed perturbation programs.

## Conclusion

We introduced **PerturbGraph**, a biologically informed graph-learning framework for predicting transcriptional responses of previously unseen gene perturbations. By representing perturbation effects as stable transcriptional programs and integrating biological priors including protein–protein interaction networks, graph-derived embeddings, transcriptional statistics, and Gene Ontology annotations, PerturbGraph enables effective propagation of perturbational information across related genes.

Across unseen perturbation benchmarks, PerturbGraph consistently outperforms classical machine learning models, perturbation-specific deep learning approaches such as scGen and CPA, and alternative graph neural network architectures. These results demonstrate that incorporating biological interaction structure provides a strong inductive bias for modeling perturbational transcriptional responses. A limitation of the current framework is that it focuses on perturbation-level transcriptional programs rather than fully modeling cell-level heterogeneity. Future work could integrate cell-type–specific regulatory networks, richer biological annotations, or generative single-cell models to further improve generalization and biological interpretability.

