## Supplementary file for "Predicting Unseen Gene Perturbation Response Using Graph Neural Networks with Biological Priors"

**6 Addtional Tables** **11**

**7 Additional Figures** **11**

**8 Reproducibility** **11**

### 1 Supplementary Overview

This supplementary document provides additional methodological details, implementation specifications, and supporting analyses for the PerturbGraph framework described in the main manuscript.

Section 5.4 presents detailed implementation information including the construction of perturbation programs, graph preprocessing steps, feature construction, and training procedures. Supplementary Table S3 summarizes the key hyperparameters used in the experiments.

In addition, we provide expanded descriptions of data preprocessing, model training settings, and evaluation procedures to facilitate full reproducibility of the results reported in the main paper.

All code used to reproduce the experiments described in this work is publicly available at:

`https://github.com/Sajib-006/PerturbGraph`

The repository contains scripts for data preprocessing, graph construction, model training, and evaluation, together with example commands for running the benchmark pipeline.

### 2 Additional Related Work

#### 2.1 Computational Modeling of Gene Perturbation Responses

Predicting gene expression responses to perturbations has been studied using a range of statistical, machine learning, and deep learning approaches. Early methods frequently relied on linear models or low-dimensional factorization techniques to capture relationships between perturbations and expression changes. These approaches are attractive due to their interpretability and simplicity but often assume limited interaction structure and may struggle to capture nonlinear or context-dependent perturbation effects [1, 2, 3, 4, 5, 6, 7, 8].

More recent work has introduced neural architectures for modeling perturbational responses, including autoencoder-based and latent variable approaches that learn compact representations of cellular expression states [9, 10, 11]. These models improve flexibility and can capture complex nonlinear mappings between inputs and perturbational outputs. However, many feature-only neural predictors treat genes independently during prediction and therefore do not explicitly leverage known molecular interaction structure.

#### 2.2 Graph-Based Learning in Functional Genomics

Graph neural networks have emerged as a powerful framework for learning from structured biological data, including protein–protein interaction networks, regulatory networks, and pathway graphs. In functional genomics, graph-based models are appealing because perturbation effects often propagate through molecular interaction networks rather than remaining localized to a single gene. By aggregating information from neighboring genes, graph neural networks can encode network context and capture dependencies that are difficult to model using independent feature vectors [12, 7, 13].

Despite these advantages, graph-based perturbation modeling presents several challenges. Prediction performance can depend strongly on graph construction, edge confidence scores, and the choice of node representation. Many existing approaches rely primarily on graph topology or a single source of biological prior knowledge. Our work builds on this line of research by pairing graph propagation with biologically enriched node features and latent perturbation program prediction.

##### 2.3 Biological Priors for Gene Representation

A large body of work in computational biology has demonstrated the value of biological priors for predictive modeling. Protein interaction networks encode functional proximity, transcription factor networks capture regulatory relationships, coexpression networks reflect shared transcriptional behavior, and ontologies such as Gene Ontology provide curated functional annotations. Pathway databases further organize genes into coordinated biological processes.

These prior knowledge sources are often used independently for feature engineering, network regularization, or downstream interpretation. In contrast, our formulation integrates multiple sources of biological context into a unified perturbation prediction framework by combining graph-derived structural embeddings, baseline transcriptional statistics, graph topology summaries, and ontology-based functional embeddings.

##### 2.4 Unseen-Perturbation Generalization

Generalization to perturbations not observed during training represents one of the most challenging and practically relevant settings in perturbation modeling. Unlike interpolation among known perturbations, this scenario requires models to infer the transcriptional impact of entirely unseen perturbation targets. This task is particularly difficult because perturbation response vectors are high-dimensional, sparse, and noisy.

Our work addresses this setting by learning in a low-dimensional perturbation program space and by using graph-based biological priors to transfer information from observed perturbations to unseen genes. This formulation differs from approaches that focus primarily on cell-level generative simulation and instead emphasizes robust perturbation-level generalization under a strict unseen-gene evaluation protocol.

#### 3 Detailed Evaluation Protocol and Baseline Definitions

##### 3.1 Detailed Evaluation Metrics

We evaluate the accuracy of predicted perturbation signatures by comparing the predicted transcriptional shift $\hat{\Delta}_i$ with the ground-truth signature $\Delta_i$ for each unseen perturbation gene $i$. Each perturbation signature is represented as a vector in gene expression space,

$$\Delta_i, \hat{\Delta}_i \in \mathbb{R}^d,$$

where $d$ denotes the number of genes in the expression representation.

**Cosine similarity.** Cosine similarity measures directional agreement between predicted and observed perturbation signatures:

$$\text{Cosine}(\Delta_i, \hat{\Delta}_i) = \frac{\Delta_i^\top \hat{\Delta}_i}{\|\Delta_i\|_2 \, \|\hat{\Delta}_i\|_2}. \tag{1}$$

A value close to 1 indicates that the predicted and observed perturbation programs are closely aligned in expression space.

**Spearman rank correlation.** We compute Spearman correlation to assess whether the relative ranking of genes by perturbation effect is preserved:

$$\rho_i = \text{Spearman}(\Delta_i, \hat{\Delta}_i). \tag{2}$$

This metric is robust to scale differences and is particularly informative when the primary goal is to recover the ordering of responsive genes.

**Precision@k for differential gene recovery.** To quantify recovery of the most strongly affected genes, we define Precision@k separately for up- and down-regulated genes. Let $U_k(\Delta_i)$ denote the set of the $k$ genes with the largest positive shifts in the observed signature and $U_k(\hat{\Delta}_i)$ the corresponding predicted set. Then

$$\text{Prec@k}_{\text{UP}} = \frac{|U_k(\Delta_i) \cap U_k(\hat{\Delta}_i)|}{k}. \tag{3}$$

Similarly, if $D_k(\Delta_i)$ and $D_k(\hat{\Delta}_i)$ denote the sets of the $k$ most negatively shifted genes in the observed and predicted signatures, then

$$\text{Prec@k}_{\text{DN}} = \frac{|D_k(\Delta_i) \cap D_k(\hat{\Delta}_i)|}{k}. \tag{4}$$

These metrics focus on the most biologically informative part of the perturbation response.

**Mean squared error.** We additionally compute mean squared error:

$$\text{MSE} = \frac{1}{d}\|\Delta_i - \hat{\Delta}_i\|_2^2. \tag{5}$$

Because transcriptomic perturbation signatures are high-dimensional and often sparse, MSE is treated as a secondary metric relative to cosine similarity and Spearman correlation.

**Aggregate evaluation.** For each metric, scores are computed independently for every test perturbation and then averaged:

$$\text{Metric}_{\text{mean}} = \frac{1}{|\mathcal{T}_{\text{test}}|} \sum_{i \in \mathcal{T}_{\text{test}}} \text{Metric}_i. \tag{6}$$

#### 3.2 Detailed Baseline Definitions

To assess the contribution of graph structure and biologically informed node features, we compare PerturbGraph against a range of baselines spanning trivial predictors, classical machine learning methods, and neural architectures.

**Mean predictor.** As a minimal reference, we use a global mean predictor that outputs the average latent perturbation program over all training perturbations. Let $\mathcal{T}_{\text{train}}$ denote the set of training perturbations and let $h_i \in \mathbb{R}^K$ denote the latent perturbation program associated with perturbation $i$. The prediction is

$$\hat{h}_{\text{mean}} = \frac{1}{|\mathcal{T}_{\text{train}}|} \sum_{i \in \mathcal{T}_{\text{train}}} h_i. \tag{7}$$

This baseline ignores perturbation-specific information and serves as a lower bound.

**k-nearest neighbors (KNN).** We next consider a nonparametric baseline that predicts the latent program of a test perturbation using the average latent programs of its nearest neighbors in feature space. Given node features $z_i$ for perturbation gene $i$, we identify its $k$ nearest training genes and predict

$$\hat{h}_i = \frac{1}{k} \sum_{j \in \mathcal{N}_k(i)} h_j, \tag{8}$$

where $\mathcal{N}_k(i)$ denotes the set of nearest training neighbors.

**Linear regression baselines.** We evaluate Ridge, Lasso, and ElasticNet regression models that map node features to latent perturbation programs. In the Ridge case, given node feature matrix $Z \in \mathbb{R}^{N \times F}$ and latent program matrix $H \in \mathbb{R}^{N \times K}$, the model learns a weight matrix $W \in \mathbb{R}^{F \times K}$ by minimizing

$$\mathcal{L}_{\text{ridge}} = \|ZW - H\|_F^2 + \lambda \|W\|_F^2. \tag{9}$$

The prediction for gene $i$ is then $\hat{h}_i = z_i W$. Lasso and ElasticNet follow the same feature-to-program framework with alternative regularization.

**Random Forest.** Random Forest provides a nonlinear, tree-based baseline that can capture interactions among node-level features without using graph propagation.

**Multilayer perceptron (MLP).** We include an MLP baseline that uses the same node features as PerturbGraph but removes graph message passing:

$$\hat{h}_i = f_{\text{MLP}}(z_i), \tag{10}$$

where $f_{\text{MLP}}$ is a feed-forward neural network. This baseline tests whether improvements arise from graph-aware propagation rather than neural capacity alone.

**Graph neural network baselines.** We compare PerturbGraph against GraphSAGE and GAT using the same interaction graph and the same node features. These baselines isolate the effect of graph neural architecture. In addition, graph-only and feature-enriched variants are used in ablation experiments to quantify the contribution of biological priors.

**Comparison protocol.** All baselines are evaluated under the same unseen-perturbation generalization setting, using identical train/validation/test gene splits and the same perturbation signature construction procedure. For methods that predict latent programs, expression shifts are reconstructed in the same low-dimensional basis. This unified protocol ensures that differences in performance reflect modeling choices rather than inconsistencies in preprocessing or split construction.

### 4 Experimental Setup Details

#### 4.1 Unseen-Perturbation Evaluation Protocol

To evaluate the ability of models to generalize to novel perturbations, we adopt an unseen-gene evaluation protocol. Let $\mathcal{G}$ denote the set of perturbation genes observed in the dataset. We partition $\mathcal{G}$ into three disjoint subsets corresponding to training, validation, and test genes:

$$\mathcal{G} = \mathcal{G}_{\text{train}} \cup \mathcal{G}_{\text{val}} \cup \mathcal{G}_{\text{test}},$$

with proportions approximately 70%, 15%, and 15%, respectively. Perturbations associated with genes in $\mathcal{G}_{\text{test}}$ are never observed during training, ensuring that evaluation measures the ability to predict transcriptional responses for previously unseen perturbations.

For each perturbation gene $i$, a pseudo-bulk perturbation signature $\Delta_i \in \mathbb{R}^d$ is constructed by averaging gene expression across cells carrying the perturbation and subtracting the mean control expression profile.

#### 4.2 Latent Perturbation Program Construction

Because perturbation signatures are high-dimensional, we project them into a lower-dimensional latent space to improve statistical stability. Let $\Delta \in \mathbb{R}^{N \times d}$ denote the matrix of perturbation signatures for training perturbations. We compute a truncated singular value decomposition (SVD):

$$\Delta \approx U_K S_K V_K^\top,$$

where $K$ denotes the latent dimensionality. The latent perturbation program for perturbation $i$ is then defined as

$$h_i = U_{K,i} S_K.$$

Models are trained to predict these latent perturbation programs, which are later mapped back to gene expression space.

#### 4.3 Interaction Graph Construction

We construct a biological interaction network using the STRING v11.5 protein–protein interaction database. Genes are represented as nodes and edges correspond to functional associations between proteins. Edge weights reflect STRING interaction confidence scores. The resulting graph is used as the underlying structure for graph neural network models.

##### 4.4 Node Feature Construction

Each gene node is associated with a feature vector integrating multiple sources of biological information:

- **Graph structural features.** Node2Vec embeddings computed on the interaction graph capture network topology and neighborhood structure.
- **Baseline transcriptional statistics.** Features derived from control cells include baseline expression level, variance, and detection frequency.
- **Graph topology statistics.** Additional structural covariates include node degree and neighborhood connectivity features.
- **Functional annotations.** Gene Ontology (GO) annotations are embedded into a compact representation using dimensionality reduction, capturing functional similarity between genes.

The final node feature vector is obtained by concatenating these feature groups.

##### 4.5 Training Details

Graph neural networks are trained to predict latent perturbation programs for genes in the training set. Model parameters are optimized using the Adam optimizer with early stopping based on validation performance. Hyperparameters including hidden dimension, learning rate, and regularization strength are selected using the validation set. The final model is evaluated on the held-out test perturbations.

#### 5 Dataset Description and Preprocessing

##### 5.1 Replogle Perturb-seq Dataset

Our primary dataset is the genome-scale Perturb-seq experiment introduced by Replogle *et al.* [14]. This dataset profiles transcriptional responses to CRISPR-based gene perturbations in human K562 cells using single-cell RNA sequencing.

After standard preprocessing, the dataset contains approximately $3.1 \times 10^5$ single cells with expression measurements for 8,563 genes. Perturbations target 1,832 genes, together with a large population of non-targeting control cells.

Each cell is associated with a guide RNA identifying the targeted gene. We construct perturbation-level signatures by aggregating single-cell measurements into pseudo-bulk profiles. For perturbation gene $i$, we compute the mean expression across all cells carrying that perturbation, and subtract the mean expression of control cells to obtain the perturbation signature:

$$\Delta_i = \bar{x}_{\text{pert},i} - \bar{x}_{\text{control}}.$$

This procedure yields a perturbation matrix

$$X \in \mathbb{R}^{N \times d},$$

where $N$ denotes the number of perturbations and $d$ the number of genes.

##### 5.2 Norman Perturb-seq Dataset

We further evaluate our method using the Perturb-seq dataset introduced by Norman *et al.* [15], which measures transcriptional responses to CRISPR-based perturbations in K562 cells.

The original dataset contains both single-gene and combinatorial perturbations. Because our goal is to evaluate prediction of single-gene perturbation responses, we filter the dataset to retain only single-gene perturbations and remove combinatorial perturbations. Perturbations with fewer than a minimum number of cells are also removed to ensure reliable pseudo-bulk estimates.

After preprocessing, the dataset contains 15,216 cells and 18 single-gene perturbations together with control cells. Gene expression is represented using a set of 2,000 highly variable genes selected during preprocessing.

Pseudo-bulk perturbation signatures are constructed using the same procedure as described for the Replogle dataset, allowing consistent evaluation across datasets.

##### 5.3 Perturbation Signature Representation

For both datasets, perturbation responses are represented as differential expression vectors relative to control cells. These signatures capture the average transcriptional shift induced by each perturbation and serve as the target outputs for perturbation prediction models.

##### 5.4 Additional implementation details

**Latent perturbation program construction via SVD.** For each perturbation, we first constructed a pseudo-bulk transcriptional response vector by averaging expression across all cells assigned to that perturbation and subtracting the mean expression of non-targeting control cells. Let $X_{\text{pert}} \in \mathbb{R}^{N \times G}$ denote the resulting perturbation-by-gene response matrix, where $N$ is the number of perturbations and $G$ is the number of genes. We then fit a truncated singular value decomposition (TruncatedSVD) *using training perturbations only* to obtain a low-dimensional latent perturbation program representation:

$$X_{\text{pert}} \approx HV^{\top},$$

where $H \in \mathbb{R}^{N \times K}$ contains perturbation-level latent programs and $V \in \mathbb{R}^{G \times K}$ contains gene loadings. In all main experiments, we used $K = 128$ latent dimensions (argument `--svd_dim 128`). The SVD basis was fit only on training perturbations and then applied to validation and test perturbations using the learned projection, preventing information leakage across splits.

**Sensitivity to latent dimension $K$.** The implementation allows the latent dimensionality to be varied through the `--svd_dim` argument. In the main reported experiments, we fixed $K = 128$, which provided a compact representation while retaining substantial variance in the perturbation response matrix. Because the current benchmark script fixes $K$ per run, sensitivity to $K$ should be interpreted based on separate reruns with alternative values (e.g., $K \in \{64, 96, 128, 192, 256\}$). In the experiments reported in this work we used $K = 128$, which provided a good trade-off between compression and reconstruction fidelity of perturbation programs. The implementation allows sensitivity analysis by varying this parameter.

**Node2Vec graph embeddings.** Graph topology features were computed using Node2Vec on the interaction network. Random walks were generated on the undirected graph and optimized using the skip-gram objective. Unless otherwise stated, the embedding dimension was set to 128 with walk length 30, 100 walks per node, window size 10, and return and in–out parameters $p = 1.0$ and $q = 1.0$. These embeddings capture higher-order network proximity between perturbation genes and were concatenated with other biological features.

**STRING graph construction and preprocessing.** Protein–protein interaction edges were derived from raw STRING links and mapped from STRING protein identifiers to gene symbols using the STRING protein information table. After mapping, self-loops were removed. We retained only interactions satisfying three criteria: (i) STRING combined score greater than or equal to 700, (ii) both incident genes present in the perturbation set, and (iii) non-self interactions. Formally, if $E_{\text{raw}}$ denotes the mapped STRING edge list, the retained graph was

$$E_{\text{STRING}} = \{(g_i, g_j, s_{ij}) \in E_{\text{raw}} : s_{ij} \geq 700,\ g_i \in \mathcal{P},\ g_j \in \mathcal{P},\ g_i \neq g_j\},$$

where $\mathcal{P}$ is the set of perturbation genes and $s_{ij}$ is the STRING combined score. Duplicate undirected edges were collapsed by sorting each pair lexicographically and keeping the maximum score for each unique gene pair. Thus, the final STRING graph is undirected, deduplicated, and score-filtered.

**Treatment of isolated nodes.** Isolated nodes were *not* removed. The node list was defined as the full set of perturbation genes, and graph objects were explicitly initialized with all nodes before edges were added. Consequently, genes without retained STRING neighbors remained in the graph as isolated nodes. For such nodes, graph-topological features default to zero where applicable, and Node2Vec embeddings also default to zero if an embedding is unavailable. This design preserves the full unseen-perturbation prediction setting rather than restricting evaluation to genes with at least one retained interaction.

**GO embedding construction.** Gene Ontology (GO) features were constructed from a gene–GO annotation table containing gene identifiers and GO term identifiers. After restricting annotations to genes present in the perturbation node list, GO terms were filtered by annotation frequency: only terms annotated to at least 5 genes and at most 500 genes were retained. From the filtered annotations, we built a binary gene-by-GO membership matrix

$$M_{\text{GO}} \in \{0, 1\}^{|\mathcal{P}| \times T},$$

where $|\mathcal{P}|$ is the number of perturbation genes and $T$ is the number of retained GO terms. We then applied TruncatedSVD to this matrix to obtain a dense GO embedding of dimension 64 (argument `--go_svd_dim 64`). If the effective rank of the matrix was smaller than 64, the representation was zero-padded to maintain a fixed feature dimensionality. Finally, each GO embedding dimension was standardized across genes using `StandardScaler`.

**GO coverage across genes.** Not all genes were required to have direct GO annotations. Genes without retained GO annotations were still included in the gene-by-GO matrix as all-zero rows because the categorical gene index was defined over the full perturbation node list. As a result, every perturbation gene received a GO feature vector, but genes lacking GO support effectively contributed zero membership before SVD projection. Therefore, coverage is complete at the feature-matrix level, although annotation support may be absent for a subset of genes.

**Graph neural network architecture and training.** The perturbation program predictor was implemented as a graph neural network with two graph convolution layers followed by a multilayer perceptron prediction head. Hidden dimension was set to 256 with dropout rate 0.15. Models were trained using the AdamW optimizer with learning rate $10^{-3}$ and weight decay $10^{-4}$. Training was performed for up to 300 epochs with early stopping based on validation cosine similarity with patience 12.

**Train/validation/test splitting.** Data splitting was performed at the *perturbation-gene level*, not at the cell level, to enforce the unseen-perturbation setting. Unique perturbation genes were partitioned into train, validation, and test subsets using `train_test_split` with user-specified fractions (`--test_frac`, `--val_frac`) and a fixed random seed. In the provided benchmark script, a single split is generated per run from the specified seed. Therefore, for Replogle, the script does *not* automatically repeat train/validation/test splitting multiple times within one execution; repeated-split evaluation must be performed by rerunning the full pipeline with different seeds. The same logic applies to Norman. If multiple-seed results are reported for Norman or Replogle, they correspond to repeated independent runs of the pipeline rather than repeated splits inside a single run. All experiments were run with a fixed random seed for reproducibility.

**Software implementation.** All experiments were implemented in Python using PyTorch and PyTorch Geometric. Data preprocessing and evaluation were performed using NumPy, pandas, scikit-learn, and Scanpy.

**Code availability.** All code used to reproduce the experiments described in this work is publicly available at `https://github.com/Sajib-006/PerturbGraph`.

### 6 Addtional Tables

### 7 Additional Figures

### 8 Reproducibility

Table 1: Evaluation of perturbation direction prediction and differential expression recovery on held-out test perturbations in the Norman dataset.

| Model | DirAcc | DE AUROC @50 | DE AUPRC @50 |
|---|---|---|---|
| Lasso | 0.602 | 0.763 | 0.194 |
| Ridge | 0.598 | 0.759 | 0.187 |
| ElasticNet | 0.609 | 0.771 | 0.209 |
| KNN | 0.611 | 0.773 | 0.232 |
| Random Forest | 0.614 | 0.775 | 0.214 |
| MLP | 0.615 | 0.780 | 0.221 |
| scGen | 0.612 | 0.774 | 0.236 |
| GraphSAGE | 0.614 | 0.775 | 0.239 |
| CPA | 0.616 | 0.779 | 0.244 |
| GAT | 0.613 | 0.782 | 0.242 |
| **PerturbGraph** | **0.619** | **0.784** | **0.257** |

Table 2: Robustness of PerturbGraph across random initialization seeds on the unseen perturbation prediction task. Results report cosine similarity and Spearman correlation between predicted and observed perturbation responses.

| Seed | Cosine ↑ | Spearman ↑ |
|---|---|---|
| 0 | 0.934 | 0.809 |
| 1 | 0.941 | 0.816 |
| 2 | 0.936 | 0.811 |
| 3 | 0.943 | 0.818 |
| 4 | 0.937 | 0.812 |
| Mean | 0.938 | 0.813 |
| Std | 0.0036 | 0.0034 |

Table 3: Key implementation defaults used in the benchmark pipeline.

| Parameter | Default value |
|---|---|
| Latent SVD dimension ($K$) | 128 |
| STRING minimum score | 700 |
| Node2Vec dimension | 128 |
| GCN hidden dimension | 256 |
| Dropout | 0.15 |
| Learning rate | $10^{-3}$ |
| Weight decay | $10^{-4}$ |
| Max epochs | 300 |
| Early stopping patience | 12 |
| Train/test fraction | 0.8 / 0.2 |
| Validation fraction | 0.15 of training genes |
| GO embedding dimension | 64 |
| GO term size filter | 5 to 500 genes |
| Pathway embedding dimension | 32 |

[10] Zijia Tang, Minghao Zhou, Kai Zhang, and Qianqian Song. Scperb: Predict single-cell perturbation via style transfer-based variational autoencoder. *Journal of Advanced Research*, 2024.
